# The activity of deep cortical layers characterizes the complexity of brain responses during wakefulness following electrical stimulation

**DOI:** 10.1101/2022.07.13.499946

**Authors:** Christoph Hönigsperger, Johan F. Storm, Alessandro Arena

## Abstract

It has been suggested that the complexity of the brain is closely related to its state of consciousness. The perturbational complexity index (PCI) has been used in humans and rodents to distinguish conscious from unconscious states based on the global cortical responses (recorded by electroencephalography; EEG) to local cortical stimulation (CS). However, it has been unclear how different cortical layers respond to CS and contribute to the resulting intra- and inter-areal cortical communication and PCI. A detailed investigation of these local dynamics is needed to understand the basis for PCI.

We hypothesized that the complexity level of global cortical responses (PCI) corresponds to variations in layer-specific activity and connectivity patterns.

We investigated global cortical dynamics and layer specific activity in mice, combining cortical electrical stimulation, global EEG, and local multi-electrode, laminar recordings from layers 1-6 in somatosensory cortex, during wakefulness and general anesthesia (sevoflurane).

We found that transition from wake to sevoflurane anesthesia correlated with a drop in global and local PCI values (complexity). This was accompanied by a local decrease in neural firing rate, spike-field coherence, and long-range functional connectivity specific to deep layers (L5, L6).

Our results suggest that deep cortical layers are mechanistically important for changes in PCI, and thereby for variations in the states of consciousness.

**Highlights:**

- Anesthesia caused a reduction in the perturbational complexity index (PCI^ST^) at both global (inter-areal) and local (intra-areal, across layers) cortical scales.
- Deep cortical layers (L6 and L5), exhibited strong connectivity with remote cortical areas during wakefulness but not during anesthesia.
- Layer 6 showed the strongest modulation of spike firing and spike field coherence compared to more superficial layers during wakefulness.

## 1. Introduction

A major goal in basal and clinical neuroscience and consciousness research is to develop objective measures of states of consciousness independently of behavioral communication, thus bypassing any deficits in sensory input or motor output (Casali, Gosseries et al. 2013, Comanducci, Boly et al. 2020). A decade ago, the perturbational complexity index (PCI) was introduced and showed ability to discriminate different levels of consciousness in humans in a variety of conditions, including wakefulness, sleep, anesthesia and unresponsive coma states (Casali, Gosseries et al. 2013, Sarasso, Boly et al. 2015), The PCI method relies on direct, local stimulation of the cerebral cortex, which in awake subjects typically causes complex, widespread and long-lasting neural responses measured by electroencephalography (EEG). The resulting highly, integrated and differentiated cortical responses in awake subjects give a high PCI value. In contrast, when consciousness is lost or strongly reduced, e.g. in slow-wave sleep or anesthesia, the cortical responses become briefer, with simpler, and stereotyped yielding low PCI values (Massimini, Ferrarelli et al. 2005, Casali, Gosseries et al. 2013). Similarly, recent experiments in rats have shown that general anesthesia decreases PCI and global functional connectivity following electrical stimulation in vivo (Arena, Comolatti et al. 2021, Arena, Juel et al. 2022). In vitro studies in cortical slices, found that application of noradrenaline and a cholinergic agonist increased the PCI (D’Andola, Rebollo et al. 2018), whereas GABA antagonists reduced PCI (Barbero-Castillo, Mateos-Aparicio et al. 2021). In contrast, rat cortical cell cultures show only moderate effects of cholinergic agonists on PCI values presumably due to a lack of appropriate circuit architecture (Colombi, Nieus et al. 2021). However it is unclear which circuit elements, e.g. cortical layers are critical for generating the integrated and differentiated brain dynamics (high PCI) which characterize the conscious state.

Indeed it is unknown how the local neural activity caused by focal cortical stimulation spreads to other parts of the cortex and how layer-specific activity and connectivity are involved. Further, we asked whether the complex dynamics typical of conscious brain states can also be found and quantified by PCI more locally, within a single cortical area, across layers. A functional readout and detailed understanding of these local dynamics is desirable from a clinical perspective where local disruptions of complexity exist after focal injury such as stroke (Sarasso, D’Ambrosio et al. 2020) and may even occur after injuries and disorders associated with perceptual deficits including visual agnosia (Riddoch and Humphreys 2003, Martinaud 2017, Whitwell, Striemer et al. 2021).

In the current study, we investigated layer-specific activity, i.e. local field potential (LFP, and multi-unit activity (MUA) and global cortical dynamics, by combining cortical electrical stimulation, global EEG and local multi electrode recordings in mouse somatosensory cortex, during wakefulness and sevoflurane anesthesia.

## 2. Materials and methods

### Legal approval and animal model

Experiments, surgeries, and animal care procedures were approved by the Norwegian food safety authority (Mattilsynet; FOTS ID: 24068) in agreement with the Norwegian regulation on animal research. Five adult C57BL/6 mice (∼20-30g, older than 4 weeks) were used in the experiments and housed within the local animal facility, at the Medical Faculty, University of Oslo. The mice were housed in ventilated, environmentally enriched cages at ∼23°C, exposed to a 12h/12h light/dark cycle and had free-access to food and water.

### Surgery and animal care

Prior to surgery, the mice were deeply anesthetized using Sevoflurane 2.5-5% (Baxter) in oxygenated (>85%), humidified air followed by subcutaneous injection of (mg/ml): 0.2 Dexamethasone, 4 Butorphanol, and 0.01 Glycopyrrolate. Surgery was performed only during absence of behavioral responses to noxious stimuli and under standard sterile procedures. For surgery mice were placed in a stereotactic frame, body temperature was maintained at ∼37-38°C by a heating pad, and after removal of skin and periosteum, small craniotomies were drilled for chronic implantation of stainless steel screws (EEG/ground electrodes, 0.5-1.2mm caliber) and insulated tungsten wires (bipolar stimulation electrodes, 50µm caliber). The electrodes were inserted into craniotomies perpendicular to the cortical surface at the following coordinates relative to the bregma position (in mm; x-axis: left (“-”) or right (“+”) hemisphere; y-axis: above (“+”) or below (“-”) bregma). Stimulation electrodes: x: +0.95, y: +2.4; x: +1.4; y: +2.4, z: 0.9 (motor cortex 2, M2). EEG electrodes: x: -1, y: +0.5; x: +1, y: +0.5 (motor cortex 1/2); x: +3, y: -1 (somatosensory cortex 1, S1); x: +1, y: -2.5 (visual 2/retrosplenial/parietal cortex, V2/Rs/MPta); x: +3.5, y: -3.5 (visual cortex 1/2 lateral, V1/V2lat); Ground electrode: x: 0, y: -4.9 (cerebellum). For acute intra-laminar recordings with linear silicone probes, a craniotomy window (∼1.5 mm diameter) was drilled at x: -2, y: -0.5 (S1). The craniotomy was covered with a silicone patch (PDMS, SYLGARD 184, Dow Corning, USA) and glued with dental acrylic. A head bar was glued in the back of the skull (behind the ground screw), perpendicular to the rostral-caudal axis, for restriction of the mice in the experimental setup. Dental acrylic was applied over the exposed skull to seal wound margins and secure electrodes in place. After the surgery, Meloxicam (5) and Buprenorphine (0.1) were injected subcutaneously and the mice returned to their home cage, followed by 3 days of post-operative care and daily injection of Dexamethasone (0.2) and Buprenorphine (0.2). During the entire period we carefully monitored hydration status, natural behavior, body condition, weight, and wound healing and checked for any possible signs of distress, infection and damage of the implant.

### Cortical electrical stimulation

We stimulated the cortex by a brief electrical stimulus (200 µs) delivered through the bipolar stimulation electrode given once every 10 seconds. In every mouse, we determined the maximum stimulus current that did not evoke whisker deflection (i.e. below motor threshold) to avoid movement- and EMG contamination of the EEG recordings. To do so, we applied 200 µs long current pulses using an isolated current stimulator (HG203, Hi-Med) triggered by a pulse generator (Pulse Pal (Sanders and Kepecs 2014)). We reduced the current amplitude until we obtained no more than a single whisker movement (determined visually, using a microscope) during 20 current stimulations (i.e. accepting a type I error of 5%). The determined current was used for triggering ERPs during the subsequent tests (mean ± standard deviation: 34 ± 17µA, n=5). For recordings, we connected EEG electrodes and a 32 channel silicone probe consisting of 2 shanks with 16 channels each (NeuroNexus, A2×16-10mm-100-500-177-CM32) to two amplifier headstages (RHS Stim/Recording System, INTAN technologies). The electrophysiological signals were amplified and referenced to ground, sampled at 20-25 kHz and band pass filtered between 0.2-8000 Hz. Prior to insertion into the brain, the silicone probe was immersed for 10 min in a fluorescet dye (Vybrant™ DiI cell-labeling solution; Invitrogen; diluted 1:10 in saline). After removal of the protective silicon window from the craniotomy in S1, the probe tip was slowly lowered at a rate of ∼2µm/s into the cortex to improve the quality of neuronal recordings (Fiáth, Márton et al. 2019). The probe was inserted into the cortex at an angle of ∼60° relative to the cortical surface, reaching a depth of 1500-1700 µm, thereby spanning the entire cortical thickness.

### General anesthesia

The general anesthetic gas sevoflurane was delivered via a mask in oxygenated (>85%), humidified air (range of concentration in 5 mice: 2.65-3.1%, mice). The depth of anesthesia was set at the minimal dose that abolished motor responses to noxious stimuli, and spontaneous movements were also not seen. In parallel, we monitored anesthesia depth by watching the spontaneous EEG activity. After experiments, mice received a lethal intra peritoneal injection of sodium-pentobarbital (140mg/kg). After suppression of the corneal reflex, mice received intra-cardiac perfusion of 4°C cold PBS or lactated Ringer’s solution followed by 4% paraformaldehyde in PBS for tissue fixation. The brain was carefully removed and stored in 4% paraformaldehyde for ∼24 hours followed by Nissl staining (Supplementary material).

### Analysis

Analysis of electrophysiological data were performed in MATLAB2016a and Python 3.7. A description of the analysis pipeline is provided in the supplementary material and on github code repository (https://github.com/chrihoni/University-of-Oslo_PCI_Mouse_Analysis).

## 3. Results

We simultaneously measured EEG and LFP responses in head-restrained mice, using two complementary methods: 1) screw electrodes, which recorded surface (epidural) EEG from frontal, parietal and occipital areas, and 2) linear silicone probes, which were inserted into the primary somatosensory (S1) cortex to record laminar LFPs in cortical layers 1-6 (Fig. 1A, B). To evoke ERPs, a bipolar stimulating electrode was chronically implanted into M2 cortex (in or near layer 5). During quite wakefulness, the spontaneous EEG was characterized by sustained low amplitude and high frequency activity (>20 Hz, Fig. 1C top), but the EEG shifted to high amplitude and low frequency activity during deep sevoflurane anesthesia that induced behavioral unresponsiveness (Fig. 1C, middle). In addition to slow-wave activity, the anesthesia also induced periods of burst-suppression activity pattern, exhibiting broadband power suppression followed by transient periods of high power activity (Fig. 1C, bottom). We found that the fraction of time spent in burst suppression was 62 ± 10 % (mean across 5 mice), quantified by the burst-suppression ratio (Vijn and Sneyd 1998, Arena, Lamanna et al. 2017). To better quantify the brain states during wakefulness and anesthesia we calculated the spectral exponent in the high frequency range (HF, 20-40 Hz), a marker of arousal level based on the decay-rate of the spontaneous EEG power spectrum in log-log coordinates (Gao, Peterson et al. 2017, Colombo, Napolitani et al. 2019, Lendner, Helfrich et al. 2020, Arena, Comolatti et al. 2021). Comparison of the periodograms showed a steeper slope of the 20-40 Hz power decay during sevoflurane anesthesia relative to wakefulness (Fig. 1D), corresponding to a decrease in the spectral exponent in all the tested mice (Fig. 1E, n = 5; *p = 0.03).

**Fig. 1:**
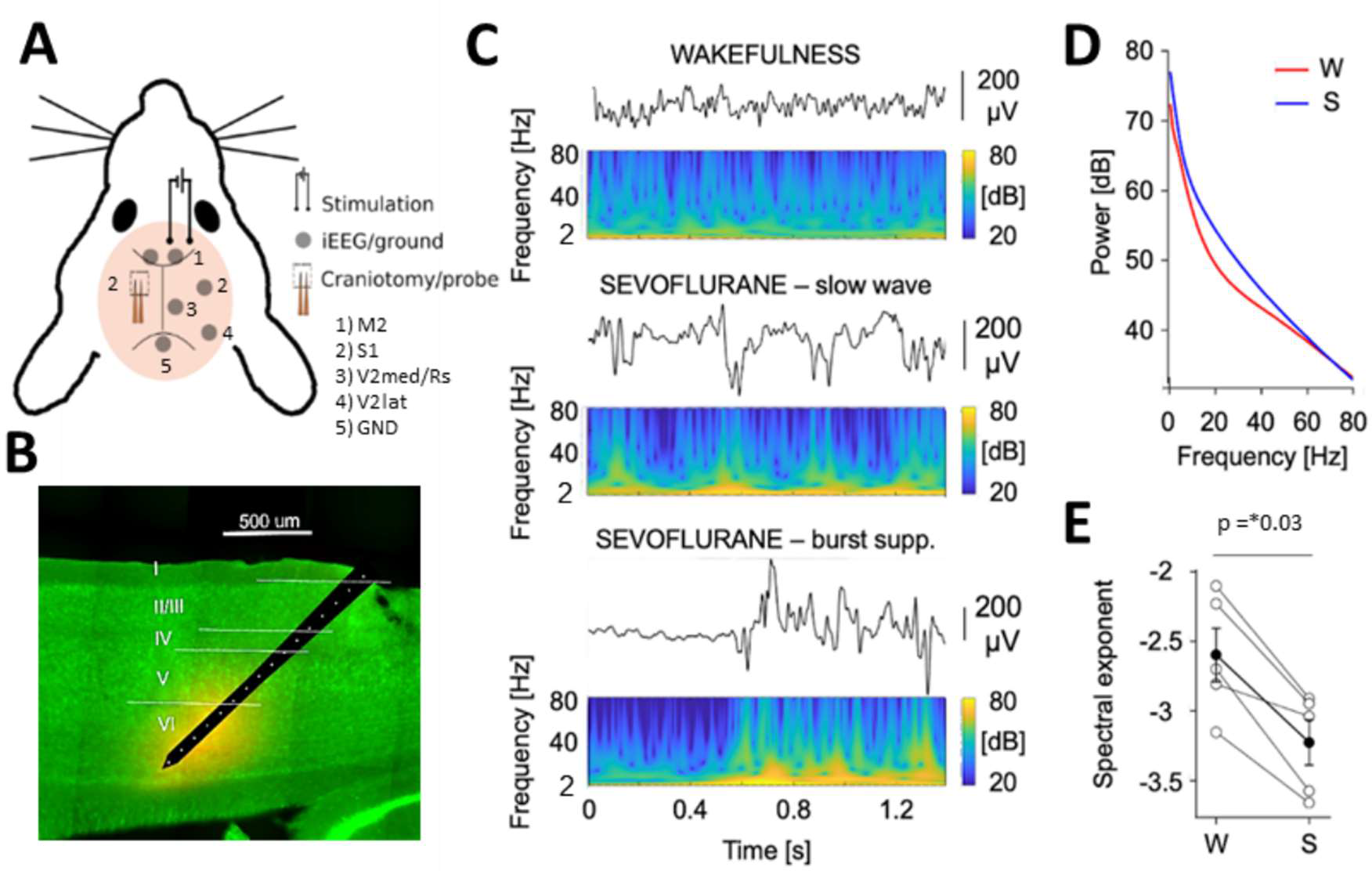
Experimental overview and spontaneous EEG activity during wakefulness and anesthetized state (sevoflurane) in mice. **A)** Experimental overview, showing placement of bipolar stimulation electrodes (M2, right hemisphere), intracranial/epidural EEG (iEEG) and ground screw electrodes (left hemisphere: M2, right hemisphere: M2, S1, V2med/Rs, V2lat), and craniotomy in S1 for placement of linear electrode array (silicone probe). **B)** Reconstructed image of silicone probe inserted into S1 cortex spanning layer 1 to 6 (sagittal view). **C)** Example traces of spontaneous EEG activity from one mouse during wakefulness (top) and sevoflurane anesthesia (middle and bottom panels). Corresponding spectrograms showing EEG power (dB) plotted vs time for frequencies ranging from 2 to 80 Hz. Note that the sevoflurane state contained a mix of both 1) slow wave activity (middle panel) consisting of increased low frequency power (<4 Hz) and nested high frequency power and 2) burst-suppression activity (bottom panel) consisting of long periods of power suppression interrupted by periods of transient high amplitude and broadband activity. The averaged “burst-suppression ratio” across five mice from electrophysiological recordings during sevoflurane anesthesia was 61.97 ± 10.07 %. **D)** Mean periodogram from 86 epochs of 2 seconds from the same mouse during wakefulness (W) and sevoflurane anesthesia (S). **E)** The spectral exponent dropped from wakefulness to sevoflurane anesthesia, highlighting a redistribution of power towards slow frequencies (n = 5 mice: wake: -2.6 ± 0.4, sevo: -3.2 ± 0.4. Mean difference in spectral exponents: -0.6, p = 0.03*. Permutation hypothesis test. Test statistic: mean difference of paired observations.

### Electrical stimulation evoked phase-locked responses across cortical areas and layers

We triggered event-related potentials (ERPs) with a single, brief electrical stimulus (200 µs), given every 10 seconds, in mouse M2 cortex (∼ layer 5). During wakefulness, the ERPs recorded by surface EEG were characterized by a sequence of voltage polarity changes over several hundred milliseconds (Fig. 2A, top). The EEG responses exhibited an increase in high-frequency power (HF, 20-80 Hz) soon after the stimulus onset, followed by a second peak of HF power at ∼160 ms (mean, n = 5 mice). The increased HF activity was still detectable up to 290 ms after stimulation and was oberserved over distant areas relative to the electrical stimulation site, ranging from M2 to the lateral V2 area (Fig. 2A, middle). We addressed the duration of the deterministic effect that the electrical stimulation had on the neuronal activity, by computing phase locking across ERPs, using the ITPC measure (David, Kilner et al. 2006, Cohen 2014, Arena, Comolatti et al. 2021). During wakefulness, stimulation triggered long lasting, phase-locked responses, lasting up to 230 ms, across several cortical areas (Fig. 2A, bottom). In contrast, during sevoflurane anesthesia, the evoked response exhibited only a brief increase in HF power and phase-locking, which quickly decayed after the stimulation (Fig. 2B).

**Fig.2:**
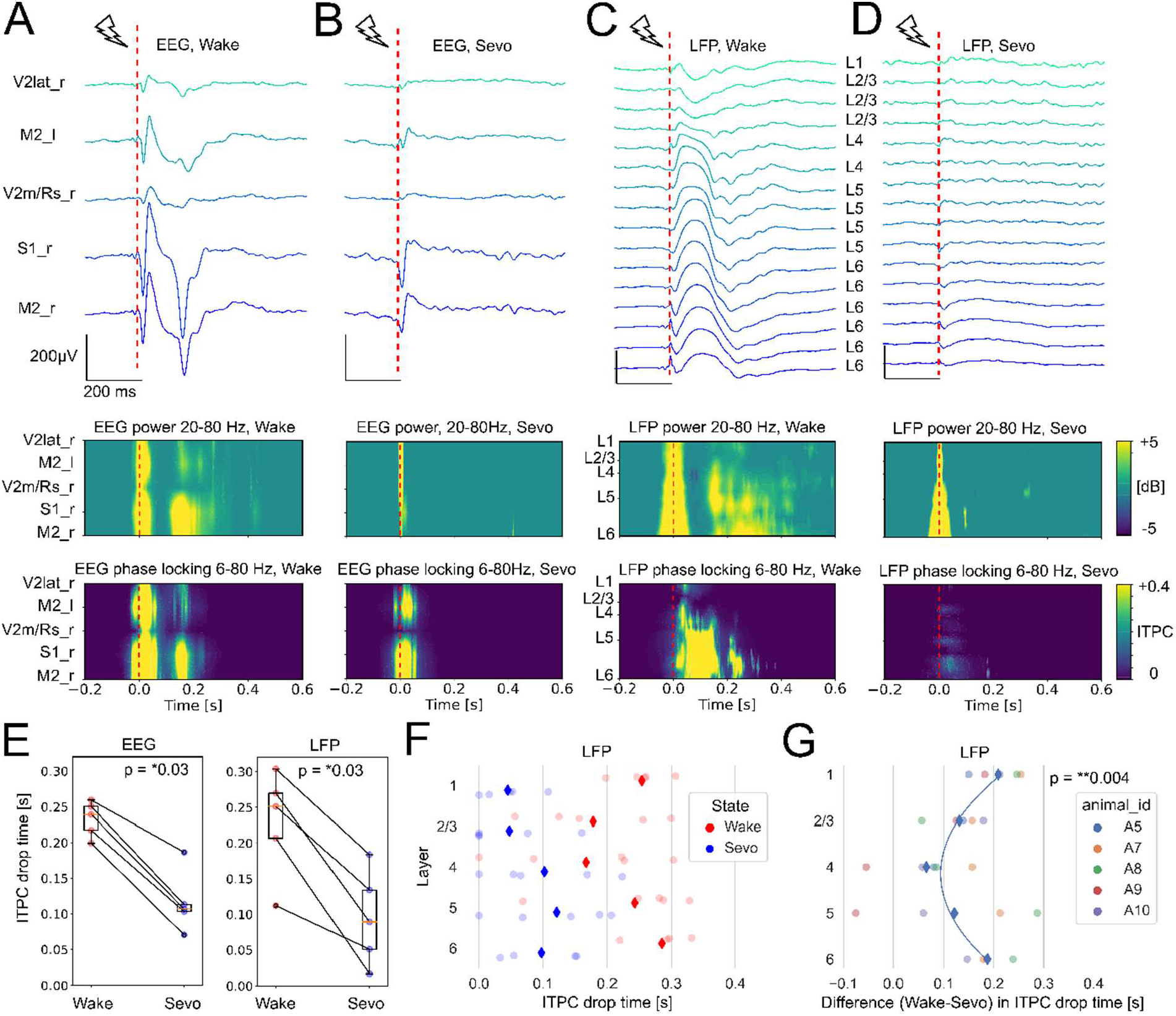
ERPs induced by electrical stimulation exhibited lower power and less phase locking, during sevoflurane anesthesia compared to wakefulness measured across cortical areas (EEG) and cortical layers (LFP in S1) **A-D**) Example traces (top) of EEG and LFP responses averaged over trials during wakefulness and sevoflurane anesthesia from one mouse. Dashed red line indicates time of electrical stimulation (in the right M2). Power spectrograms (middle) and ITPC phase-locking plots (bottom) of EEG and LFP averaged over all trials during wakefulness and sevoflurane anesthesia. **E, left**) ITPC drop time (in seconds) of EEG signals averaged across cortical areas in seconds (n = 5 mice): wake (0.23 ± 0.02 s), sevo (0.12 ± 0.04 s). Mean difference in ITPC drop time (s): -0.12, *p = 0.03). **E, right**) ITPC drop time (s) of LFP signals averaged across cortical layers: wake (0.23 ± 0.07), sevo (0.1 ± 0.07). Mean difference in ITPC drop time (s): -0.13, p = *0.03. Permutation hypothesis test. Test statistic: mean difference of paired observations. **F)** Mean LFP ITPC drop time (s) from 6-80Hz in each cortical layer during wakefulness (red diamonds; L1: 0.25, L2/3: 0.18, L4: 0.17, L5: 0.24, L6 0.28; n = 5), and during sevoflurane anesthesia (blue diamonds; L1: 0.04, L2/3: 0.05, L4: 0.1, L5: 0.12, L6: 0.1, n = 5). **G)** Difference in mean LFP ITPC drop time (s) between wakefulness and sevoflurane: (blue diamonds, L1: 0.21, L2/3: 0.13, L4: 0.07, L5: 0.12, L6 0.19; n = 5 mice, A5 - A10). The mean difference in the ITPC drop time across layers showed a C-shaped profile when fitted with 2^nd^ order polynomial. R^2^ adjusted = 0.93, p = **0.004. Permutation hypothesis test. Test statistic: 2^nd^ order coefficient of fitted polynomial. (Null hypothesis: The ITPC drop follows a uniform profile across layers.).

We next investigated if the global changes in ERP power and phase coherence are also present at a smaller scale and thus detectable across superficial and/or deep layers. To do so we inserted a linear electrode array into the S1 cortex and recorded laminar LFP responses across the cortical layers. During wakefulness, ERPs were detectable in all cortical layers, which gradually changed voltage dynamics between the different recording positions (Fig. 2C, top). About 100 ms after the stimulation, layers 4, 5 and 6 exhibited a slow, positive wave followed by a series of faster voltage deflections (Fig. 2C, top). The fast deflections were characterized by an increase in HF power, most prominent in layers 5 and 6 (Fig. 2C, middle). Likewise, phase-locking was more pronounced and long-lasting in the deep layers, especially in layer 6, lasting up to 300 ms after the stimulation (Fig. 2C, bottom). During sevoflurane anesthesia the ERPs were substantially reduced (Fig. 2D, top), and characterized by short-lasting HF power and phase-locking activity after the stimulus (Fig. 2D, middle, bottom). We found that the ITPC drop time was similar between the EEG and LFP responses, and both of these measures decreased from wakefulness to sevoflurane anesthesia (Fig. 2E, p = *0.03). Interestingly, during wakefulness the cortical layer with the latest ITPC drop time was layer 6 with ∼300 ms, followed by layer 1 and layer 5 (Fig. 2F, red diamonds), whereas all layers showed a reduced ITPC drop time during sevoflurane anesthesia (Fig. 2F, blue diamonds). To address which cortical layer is most affected by anesthesia, we calculated the mean difference in ITPC drop time between wakefulness and anesthesia (Fig. 2G). Here, layers 1 and 6 showed the largest difference in ITPC drop time, and layer 4 the smallest difference across states (Fig. 2G, blue diamonds). Thus, the duration of phase-locking after electrical stimulation was not uniform across the cortical layers, and the difference in mean LFP ITPC drop time between wakefulness and anesthesia appeared to have a “parabolic” or “C-shaped” profile, reaching its maxima in the most superficial (L1) and deepest (L6) cortical layers (Fig. 2G, p = **0.004).

### Perturbational complexity at global and local scales correlate during wakefulness and anesthesia

Next, we quantified the spatiotemporal complexity of ERPs associated with EEG and LFP voltage responses, using PCI^st^ (Comolatti, Pigorini et al. 2019). We first calculated the time course of PCI^st^ derived from EEG signals across cortical areas (Fig. 3A), within overlapping time windows of 100 ms (Arena, Comolatti et al. 2021). Soon after stimulation, i.e. within the first ∼50 ms, the magnitude of PCI^st^ was similar during wakefulness and anesthesia, but diverged within the next ∼200-300 ms (Fig. 3A). The PCI^st^ derived from LFP signals in cortical layers showed a similar time course (Fig. 3B). We determined the time period when PCI^st^ of both EEG and LFP diverged between the wakefulness and anesthesia state by calculating the difference (Δ) in 95% confidence intervals (CI) of PCI^st^ (Fig. 3C, black horizontal lines). This time period lasted from 40 to 320 ms, and within this window the PCI^st^ decreased from wake to anesthesia for EEG (Fig. 3D, p = *0.03) and LFP responses (Fig. 3E, p = *0.03). Notably, within a time window of ∼50-100 ms the difference in 95% CI was slightly larger and non-overlapping in EEG relative to LFP responses, (Fig. 3C). This finding might suggest a higher sensitivity of EEG signals to detect early state-dependent changes in PCI^st^. In conclusion, we found that complexity of global EEG and local LFP responses positively correlate between wakefulness and anesthesia state (Fig. 3F, R^2^ = 0.47, p = *0.018).

**Fig. 3.**
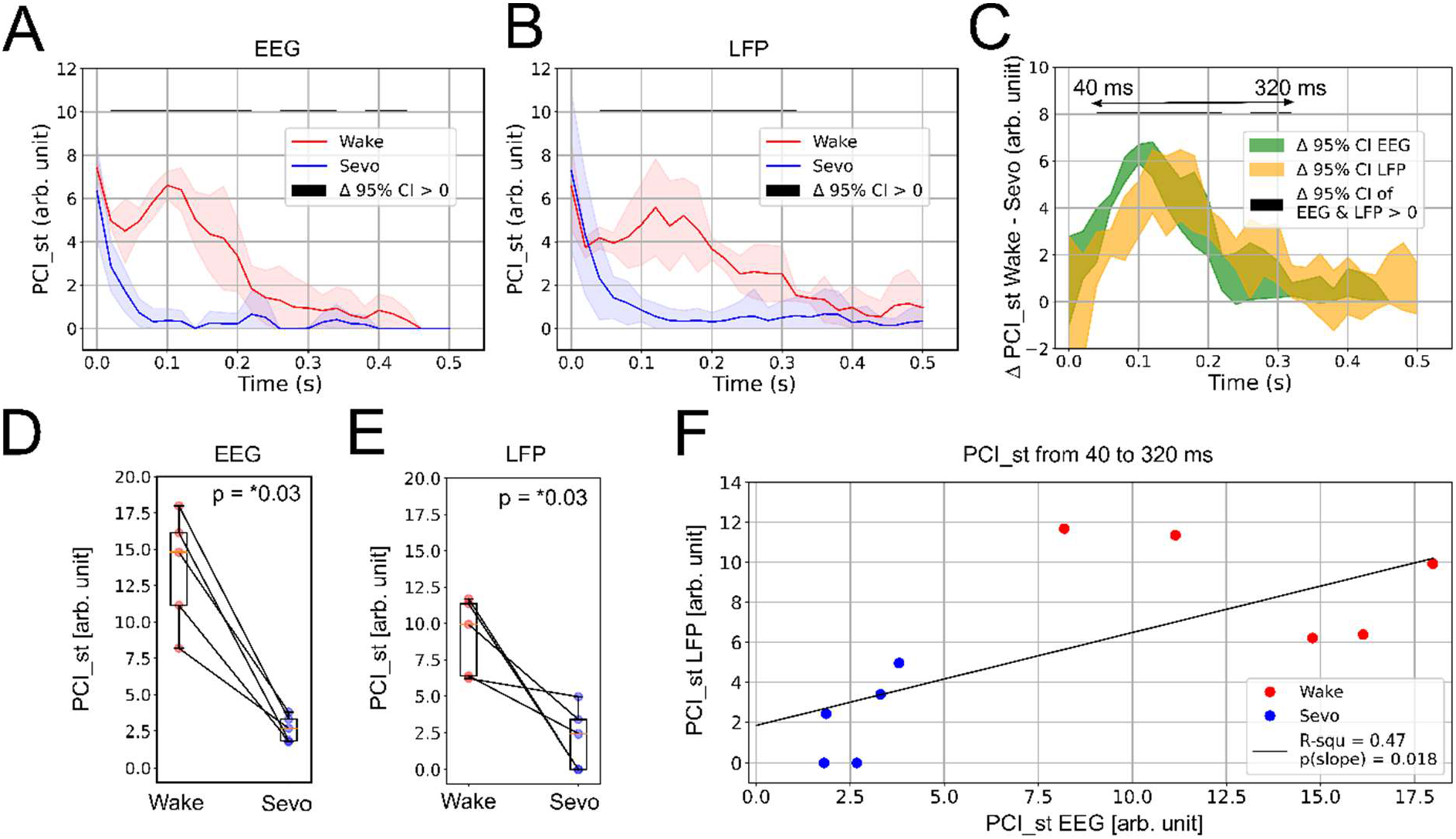
PCI^st^ of EEG and LFP responses correlate during wakefulness and sevoflurane anesthesia. **A-B)** Time course of PCI^st^ calculated from mean EEG (A) and LFP (B) responses during wakefulness and sevoflurane anesthesia (Mean in bold, std in shades). Black horizontal line indicates time points where mean PCI^st^ values are higher in wakefulness compared to sevoflurane condition, by calculating the difference of 95% confidence intervals of mean PCI^st^ values during wakefulness and sevoflurane anesthesia (Δ > 0). **C)** Overlay of EEG (green) and LFP (orange) PCI^st^ time courses showing difference in 95% confidence intervals of mean PCI^st^ values (wakefulness – sevoflurane, shaded areas). Black horizontal line indicates time points where mean PCI^st^ values of both EEG and LFP are higher during wakefulness compared to sevoflurane condition (Δ > 0), i.e. within a range of 40 to 320 ms. **D)** PCI^st^ calculated from EEG responses between 40 to 320 ms (range of horizontal lines in C): wake (13.65 ± 3.95), sevo (2.69 ± 0.88). Mean difference in PCI^st^ (wake-sevo): 10.96, p = *0.03. Permutation hypothesis test. Test statistic: mean difference of paired observations. **E)** PCI^st^ calculated from LFP responses between 40 to 320 ms: wake (9.11 ± 2.65), sevo (2.16 ± 2.17). Mean difference in PCI^st^ (wake-sevo): 6.96, p = *0.03. **F)** PCI^st^ of EEG and LFP correlate positively during wakefulness and anesthetized state. R^2^: 0.47; slope: 0.47; p (slope) = *0.02. Permutation hypothesis test. Test statistic: slope of linear regression. Null hypothesis: slope = 0.

### Layer 6 shows strongest increase in spike firing and spike-field coherence following electrical stimulation

Inter-areal coherence can emerge without local spiking entrainment due to afferent synaptic inputs (Schneider, Broggini et al. 2021). Hence, we asked whether spatiotemporal complexity of EEG and LFP responses (Fig. 2, 3) might result from 1) spiking only in afferent fibers evoked by the electrical stimulation, which project to various cortical and laminar targets, or 2) result from afferent synaptic activity causing modulation of spike firing in other cortical areas. To address this, we analyzed the multi-unit spike activity (MUA) of high-pass filtered LFP responses in S1 cortex (Fig. 4). During wakefulness, the spontaneous MUA spike rate markedly differed between cortical layers (Fig. 4A) with layer 5 showing the highest and layer 6 the lowest average spike rates during baseline time periods (spikes/s, l6: 10 ± 12, l5: 62 ± 51, l4: 57 ± 46, l2/3: 19 ± 27). Immediately after the electrical stimulation, the MUA spike rate dropped, causing a period of neuronal silence, (“off-period”, Fig. 4B), which lasted ∼90 milliseconds in average across layers and animals. Interestingly, the off-period was followed by an increase in MUA spike rate (“on-period”, see Methods), during which layer 6 showed the largest relative increase in spike firing compared to more superficial layers in wakefulness condition (Fig. 4B, C). Next, we investigated the MUA responses during sevoflurane anesthesia. Here, the average MUA activity was sparse in all cortical layers and alternated between periods of spike bursting and silence (Fig. 4D). Further, we did not observe a consistent increase in MUA rate relative to baseline between deep or superficial layers (Fig. 4E, F), when compared to the wakefulness on-period. Besides, the MUA off-period showed a higher variability during anesthesia, being absent in 4/5 mice (one of these mice shown in Fig. 4E, F), and present in 1/5 mice in layer 5 (Supplementary Fig. 1). To summarize our findings, we quantified the MUA firing rate within the on-period across all animals (Fig. 4G, I). During wakefulness, layer 6 showed a larger increase in firing rate compared to layer 5, 4 and 2/3 (Fig, 4G). In contrast, during sevoflurane anesthesia the change in firing rate was not significantly different in layer 6 compared to layer 5, 4 and 2/3 (Fig. 4I). We also compared the impact of anesthesia on MUA activity in each layer and found that layer 4-6 exhibited reduced firing rates in anesthesia vs. wakefulness condition (l2/3: -0.17, p=0.09; l4: - 0.25, p=*0.02; l5: -0.23, p=***4e-4; l6: -0.51, p=*0.02; Permutation hypothesis test. Test statistic: Mean difference in normalized MUA rate of paired observations, i.e. sevo-wake). Finally, we asked how consistent spike firing is relative to the phase of the ongoing LFP and whether the synchronization between the LFP and MUA spike times differs across cortical layers by calculating the spike-field coherence (SFC). As SFC depends on the firing rate (Lepage, Kramer et al. 2011) we plotted SFC as a function of mean MUA rate for each channel during the on-period (Fig. 4H, J) and quantified the relationship of SFC and spike rate using a linear model. Here, we determined SFC at theta frequencies (6-8Hz), which exhibited the longest phase locking of the LFP in wake mice (range: 100 to 300 ms, n = 5) relative to higher frequency bands (alpha, beta, gamma). In wake mice we found that layer 6 exhibited the best fit of the linear model compared to more superficial layers (Fig. 4H, R^2^ = 0.54, p = ***3e-5). Hence, layer 6 fired more consistently relative to the LFP at theta frequencies, if compared to e.g. layer 5 which exhibited both high and low coherence values independent of the underlying spike rate (Fig. 4H, R^2^ = 0.05, p = 0.09). Unexpectedly, layer 4 exhibited lower coherence values at higher spike rates, thereby exhibiting an opposite trend as compared to other layers (Fig. 4H, R^2^ = 0.35, p = *0.019). Similarly to layer 6, the SFC of layer 2/3 showed a significant positive relation with the firing rate, even with lower R^2^ (Fig. 4H, R^2^ = 0.27, p = **0.005). Moreover, despite similar intercepts, the slope of the model was higher for layer 6 compared to layer 2/3 (L6, intercept = 0.14, slope = 0.006; L2/3, intercept = 0.17, slope = 0.0025). This indicates that the SFC of layer 6 became quickly the highest, with higher spike rate. In anesthetized mice, we found that layer 6 did not show an increase in MUA rate during the on-period (Fig 4I) or increase in coherence as a function of spike rate (Fig. 4J) relative to more superficial layers. In addition to coherence at theta frequencies we also quantified coherence at alpha, beta and gamma frequencies (Supplementary Figure 2). Notably, in wake mice coherence values dropped from low to high frequency bands across different layers, suggesting that lower frequency bands enable a more effective spike-field coupling following cortical stimulation.

**Fig. 4:**
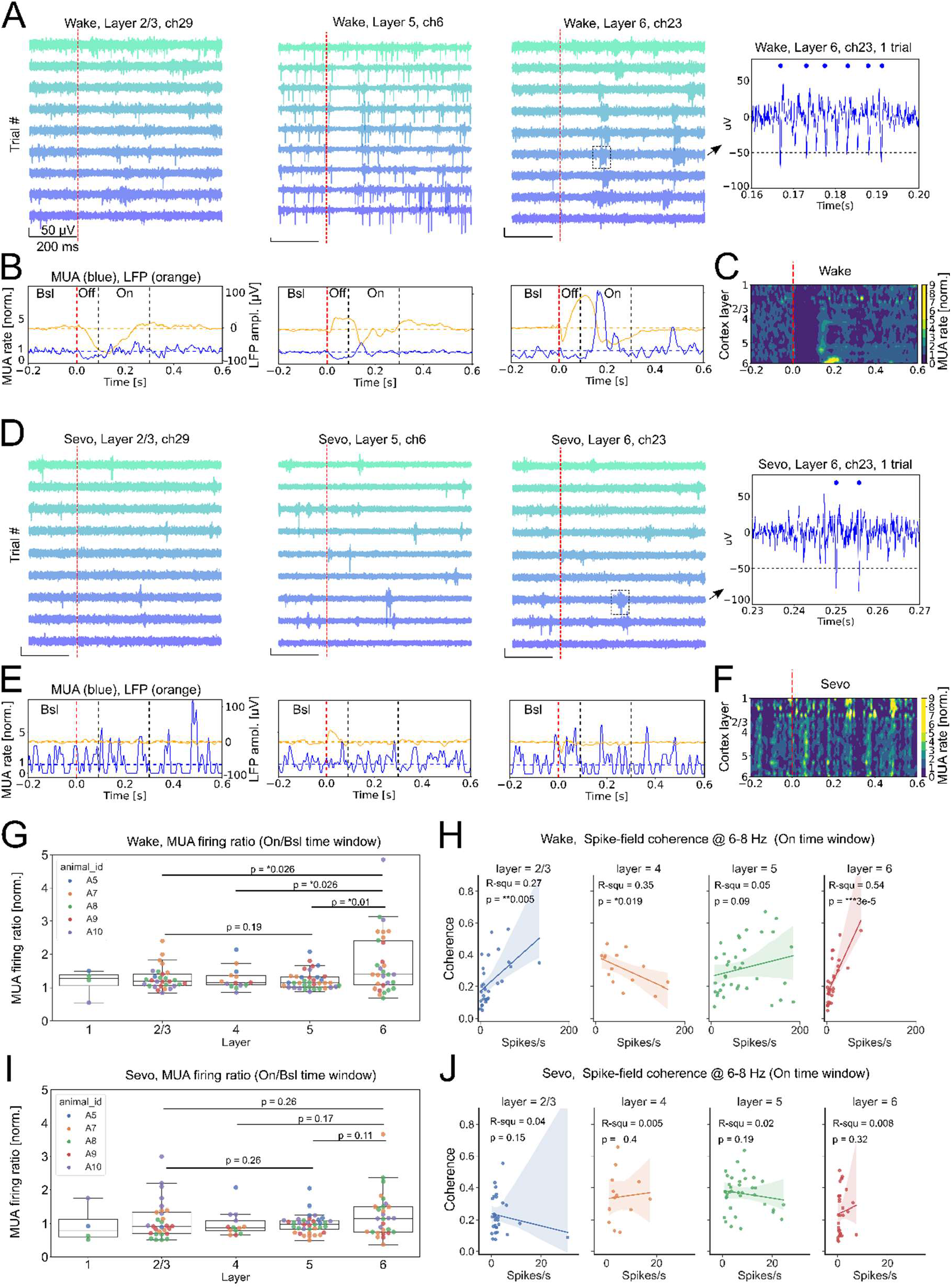
Layer 6 shows stronger increase in multi-unit spike activity (MUA) and spike-field coherence following electrical stimulation compared to other cortical layers. **A)** Examples of band pass filtered MUA traces recorded over consecutive trials in one mouse during wakefulness, from 3 different channels located in layer 2/3 (left), layer 5 (middle), and layer 6 (right). Vertical red lines indicate time of electrical stimulation (in M2, see Fig. 1A). Note close-up of layer 6 MUA activity during one trial (far right). Horizontal line indicates threshold for MUA detection (-50uV). Detected spikes are marked with blue dots. **B)** Overlay of mean MUA rate (blue, averaged across 118 trials and normalized to baseline) and mean LFP responses (orange, uV) during wakefulness in layer 2/3, 5 and 6 (from left to right, same channels as in A). Note that the baseline (bsl) is followed by an off period (reduction in MUA firing) and on period (increase in MUA firing) after electrical stimulation. **C)** Heat map of mean MUA rate, showing firing activity of all cortical layers and channels relative to baseline in one mouse. Note the strong increase in MUA firing in layer 6, similar to plots shown in A, B, right. **D-F)** Example traces and mean MUA rates during sevoflurane anesthesia. Note sparse but synchronized MUA activity across channels, and lack of consistent off/on period following electrical stimulation. **G)** Summary of mean MUA firing rates during wakefulness across cortical layers (“on” time window, relative to baseline, n = 5 mice (A5-A10); dots represent recording sites): l2/3 (1.3 ± 0.3), l4 (1.3 ± 0.3), l5 (1.2 ± 0.3), l6 (1.7±0.9). Layer 6 exhibits larger increase in firing rate compared to layer 2/3, 4, 5 during wakefulness (p = *0.026, *0.026, *0.01). Bootstrap hypothesis test (Test statistic: difference in means, H0: no difference in means). We corrected p-values for multiple comparison using the Holm-Bonferroni Method. **H)** Spike-field coherence calculated between LFP (6-8 Hz) and MUA during wakefulness (“on” time window, n = 5 mice). Layer 6 shows stronger correlation between coherence and MUA rate (spikes/s) compared to other cortical layers (r-squared, slope coefficient, p-value (slope_coeff)): l2/3 (0.27, 0.0025, **0.005), l4 (0.35, -0.0014, *0.019), l5 (0.05, 7e-4, 0.09), l6 (0.54, 0.006, ***3e-5). Shaded areas indicate 95% CI of fitted regression models. **I)** The mean MUA firing rates during sevoflurane anesthesia do not differ between layer 6 and other cortical layers: l2/3 (1.1 ± 0.6), l4 (1 ± 0.4), l5 (1 ± 0.3), l6 (1.2±0.7), (p = 0.26, 0.17, 0.11). **J)** Spike-field coherence during sevoflurane anesthesia indicates no correlation between coherence and MUA rate in any cortical layer: l2/3 (0.04, -0.004, 0.15), l4 (0.005, 0.002, 0.4), l5 (0.02, -0.0025, 0.19), l6 (0.008, 0.007, 0.32). Permutation hypothesis test (Test statistic: slope coefficient of linear regression, H0: no linear relationship between rate and coherence, i.e. slope =0).

### Functional connectivity between deep cortical layers and remote cortical areas is high during wakefulness and drops during anesthesia

The PCI values quantified in Figure 3 estimate integration and differentiation of a neural network depending on its state (Casali, Gosseries et al. 2013, Comolatti, Pigorini et al. 2019). Coherently, it was recently shown that high PCI values are correlated with highly connected brain areas with differentiated connectivity patterns (Arena, Comolatti et al. 2021). Hence, we asked which cortical layers exhibit the strongest connectivity and differentiation relative to remote cortical areas following electrical stimulation. To answer this question we calculated connectivity, between all LFP and EEG channel pairs using inter site phase clustering (ISPC, Fig. 5). During wakefulness, functional connectivity persists over a time scale of >200ms, in the frequency range of 5-20Hz, between deep cortical layers and contralateral, distant cortical areas (Fig. 5A; e.g. EEG right M2, with LFP left S1). By comparing the mean ISPC across layers in wake mice, we found that layer 5 and 6 exhibited larger connectivity with the opposite hemisphere compared to layer 2/3, while layer 6 showed also larger connectivity than layer 1 (Fig. 5B-D, p = *0.04, p < ***1e-5, p < ***1e-5). Conversely, in the anesthetized mouse, the connectivity of the same channel pairs is abolished or strongly reduced (Fig. 5A) and we did not detect layer-dependent differences in averaged connectivity (Fig. 5B-D). Overall, the functional connectivity calculated by the mean ISPC across all recording sites was higher in wake compared to anesthetized mice (Fig. 5E, p = *0.03).

**Fig.5:**
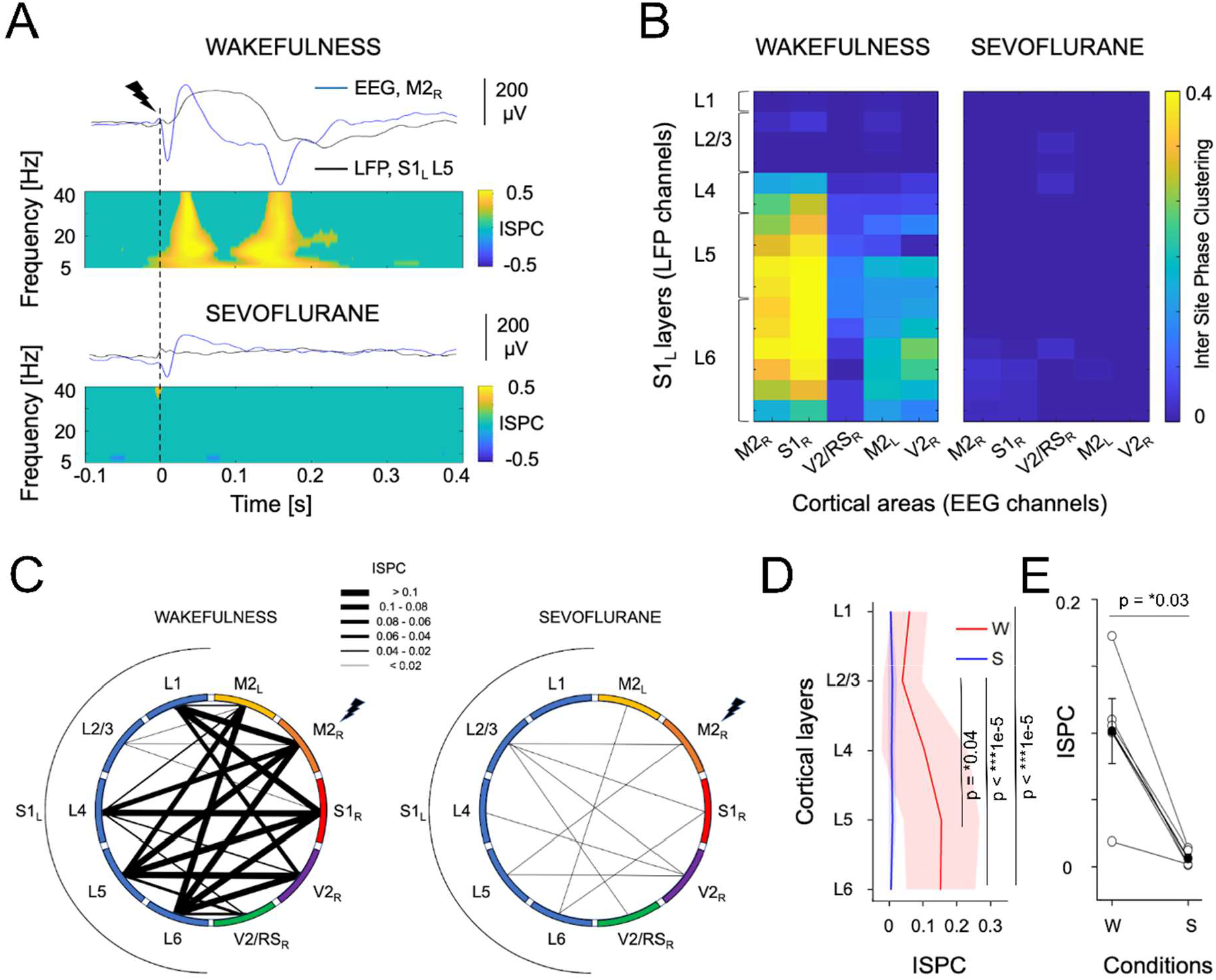
Functional connectivity between deep cortical layers and remote cortical areas is abolished during anesthesia. **A)** Example of epidural EEG and layer 5 LFP activity in response to single pulse electrical stimulation of the right secondary motor cortex (M2R, dashed line) from the same rat during wakefulness (top) and sevoflurane anesthesia (bottom). The mean ERPs recorded from the right secondary motor cortex (EEG, M2R) and from the left primary somatosensory cortex (LFP, S1L) are superimposed in the upper panel to highlight phase coherence. The variation of functional connectivity along time and across the reported cortical regions is quantified in the bottom panels, with the inter site phase clustering (ISPC), across trials (n=86). **B)** The connectivity matrixes report the mean positive ISPC values between 90 and 300 ms after the stimulus onset and in frequency range 6-8 Hz, across cortical areas (EEG channels) and cortical layers of S1L(LFP channels), from the same mouse and conditions of A). The median ISPC values (n = 5) are coded as line thickness and reported in **C).** **D)** The mean ISPC ranging from 90-300 ms calculated at 6-8 Hz across conditions, for each layer, after averaging across corresponding channels, cortical areas and mice (N=5, left). During wakefulness, the ISPC is differentiated across cortical layers (principal effect of layers, Friedman test; p = *0.03), with higher values in deeper layers. During sevoflurane anesthesia ISPC was more homogenous and no principal effect of layers could be detected (Friedman test; *P* = 0.77). During wakefulness the mean ISPC is larger in layer 6 compared to superficial layers 1 and 2/3 (p < ***1e-5) and larger in layer 5 compared to layer 2/3 (p = *0.04). During anesthesia, no mean ISPC difference is found between layers (p = 0.5). Bootstrap hypothesis test (Test statistic: difference in means, H0: no difference in means). We corrected p-values for multiple comparison using the Holm-Bonferroni Method. We found no mean difference in ISPC between wakefulness and sevoflurane condition within superficial and deep layers, when correcting for multiple comparisons using Holm-Bonferroni method. L1, p = 0.18; L2/3, p = 0.18; L4, p = 0.18; L5, p = 0.15; L6, p = 0.15. On the contrary the ISPC during wakefulness was higher than during anesthesia in deeper layers when uncorrected for multiple comparisons (L5, p = *0.03; L6, p = *0.03). Permutation hypothesis test. Test statistic: mean difference of paired observations. **E)** Overall, the functional connectivity quantified by the mean ISPC between cortical layers of S1L and other five cortical areas was always higher during wakefulness and dropped down with sevoflurane anesthesia (wakefulness: 0.1 ± 0.06 (STD), sevoflurane: 0.006 ± 0.006 p = *0.03). Permutation hypothesis test. Test statistic: mean difference of paired observations.

## 4. Discussion

### Aim of the study

PCI was originally developed, tested, and used to quantify the joint differentiation and integration (complexity) of electrophysiological responses across large parts of the human cerebral cortex, following a local cortical perturbation, e.g. by a local TMS or electrical stimulation (Casali, Gosseries et al. 2013, Comolatti, Pigorini et al. 2019). However, it is unclear if similar complex dynamics are also present within a single cortical area in intact brains, with integrated and differentiated neuronal activations across superficial and deep layers. Besides, it is also uncertain if layer-specific connectivity and causal firing of local neurons are involved in sustaining the complex dynamics across cortical areas, seen at the EEG level. To study global and local complexity in concert, we combined electrical stimulation in mouse cortex with EEG and multisite LFP recordings in layer 1-6, and analyzed PCI, connectivity and local spike activity during wakefulness and sevoflurane general anesthesia.

### Global and local PCI values

We observed that general anesthesia caused a drop in both global (EEG-based) and local (LFP-based) PCI values, which is plausible for a brain-wide effect of a systemically applied anesthetic. A previous study using EEG in humans showed that global and local PCI measures could also diverge, during loss of local but not global signal complexity following focal brain injury (Sarasso, D’Ambrosio et al. 2020). While a local measure of complexity may not always predict the overall state of a system, it certainly helps to understand how integrated a cortical column is with the rest of the brain. In our study, global and local PCI values co-vary, arguing that S1 is part of an integrated brain response, where layer-dependent effects can be studied more in detail.

### Layer-specific LFP dynamics

In wake mice (high PCI), we found that ERPs were detectable in all cortical layers, which gradually changed voltage dynamics between the different recording positions. This suggests that ERPs were not due to afferent synaptic activity restricted to superficial or deep layers alone. However, a straightforward conclusion is difficult as single cells can also exhibit sinks and sources in different layers depending on the synapses being activated (Herreras 2016). To minimize the contribution of remote sources to the field potential, we referenced LFPs pairwise between electrodes at the same cortical depth (Methods). Interestingly, the duration of the phase-locked ERP, quantified as ‘ITPC drop time’ exhibited the longest phase locking in layers 6, 5, 1, and shortest duration in layer 2/3 and 4 in wake mice. The short-lasting response in layer 4 may be related to the site of our electrical stimulation in deep M2 cortex, as layer 4 is the main direct target of sensory inputs from the thalamus (Harris and Shepherd 2015), and hence may not receive strong inputs through “higher-order” M2 projections. Conversely, long lasting phase-locked responses in layers 6, 5, 1 match more closely with thalamo-cortical inputs from matrix- and intralaminar-type projections (Harris, Mihalas et al. 2019). Notably, stimulation of intralaminar nuclei increase cortical arousal in humans (Schiff, Giacino et al. 2007), monkeys (Redinbaugh, Phillips et al. 2020, Bastos, Donoghue et al. 2021) and rats (Xu, Galardi et al. 2020), however the role of subcortical nuclei in sustaining complex activity following cortical stimulation remains to be shown.

### MUA activity and functional connectivity in layer 5, 6

Interestingly, we found that layer 6 showed the largest increase in MUA firing rate and most consistent theta-band spike-field coherence relative to more superficial layers in wake mice. Conversely, during anesthesia we did not detect an increase in firing rate in any cortical layer. In addition, the spike-field coherence was not related to the ongoing firing rate anymore. Our finding that layer 6 showed a strong increase in firing rate is supported by a recent study which found that within layer 6, layer 6b receives the strongest input from monosynaptic long-range layer 5/6 projections from contralateral S1 and motor cortex (Zolnik, Ledderose et al. 2020), which are close to our stimulation site in deep contralateral M2 cortex. Indeed, our data show that functional connectivity is strongest between electrodes located in layer 6/5 and contralateral S1/M2 cortex in wake mice. Thus, the connectivity in layer 6/5 overlaps with the dendrites of layer 6 neurons which span deep layers, where long-range cortico-cortical projections traverse (Vandevelde, Duckworth et al. 1996, Watakabe and Hirokawa 2018). It is worth noting that the dendrites of other neurons overlap with long-range projections as well, and that measures of complexity likely depend on interplay between different (sub-) cortical cells interacting with layer 6/5 (Zolnik, Ledderose et al. 2020, Afrasiabi, Redinbaugh et al. 2021).

### Limitations of current study and future perspectives

The use of EEG and laminar silicone probes has the advantage of measuring neural events with both slow and high temporal resolution, but it does not provide cell type specific information related to the PCI measure. Ideally, the use of transgenic mouse lines, which express Ca2+/voltage-sensitive indicators in various cortical and subcortical cells, would give a more comprehensive picture of neural activity related to PCI. In addition, optogenetic stimulation of specific cell-types could identify their relative contribution to brain complexity compared with conventional electrical stimulation, which presumably excites a variety of cells and fibers simultaneously.

## 5. Conclusions

Our data indicate that transition from wakefulness to general anesthesia (sevoflurane) correlates with a drop in both global and local perturbational complexity, as quantified by PCI^ST^. Within a single cortical area (S1), we observed long-lasting and deterministic neuronal activations that were differentiated across layers during wakefulness and short lasting and simplified during anesthesia. Here deep layers showed the strongest causal engagement following cortical stimulation including increase in spike firing and spike-field coherence (layer 6) and long-range functional connectivity (layer 5 and 6) in wake mice. Our results suggest that deep cortical layers are mechanistically important for changes in PCI, and thereby variations in the states of consciousness.

## Abbreviations

PCI-ST: (perturbational complexity index - state transition)
CS: (cortical stimulation)
EEG: (electroencephalography)
LFP: (local field potential)
MUA: (multi-unit activity)
ERP: (event-related potential)
ITPC: (inter-trial phase clustering)
ISPC: (inter-site phase clustering)
SFC: (spike-field coherence)

## Credit authorship contribution statement

**Christoph Hönigsperger:** Data collection/curation, Formal analysis, Visualization, Writing – original draft, Experimental design

**Johan F. Storm:** Writing – Review & Editing, Funding acquisition, Experimental design

**Alessandro Arena:** Data collection/curation, Formal analysis, Visualization, Writing – Review & Editing, Experimental design

## Supplementary material

**Supplementary Fig. 1:**
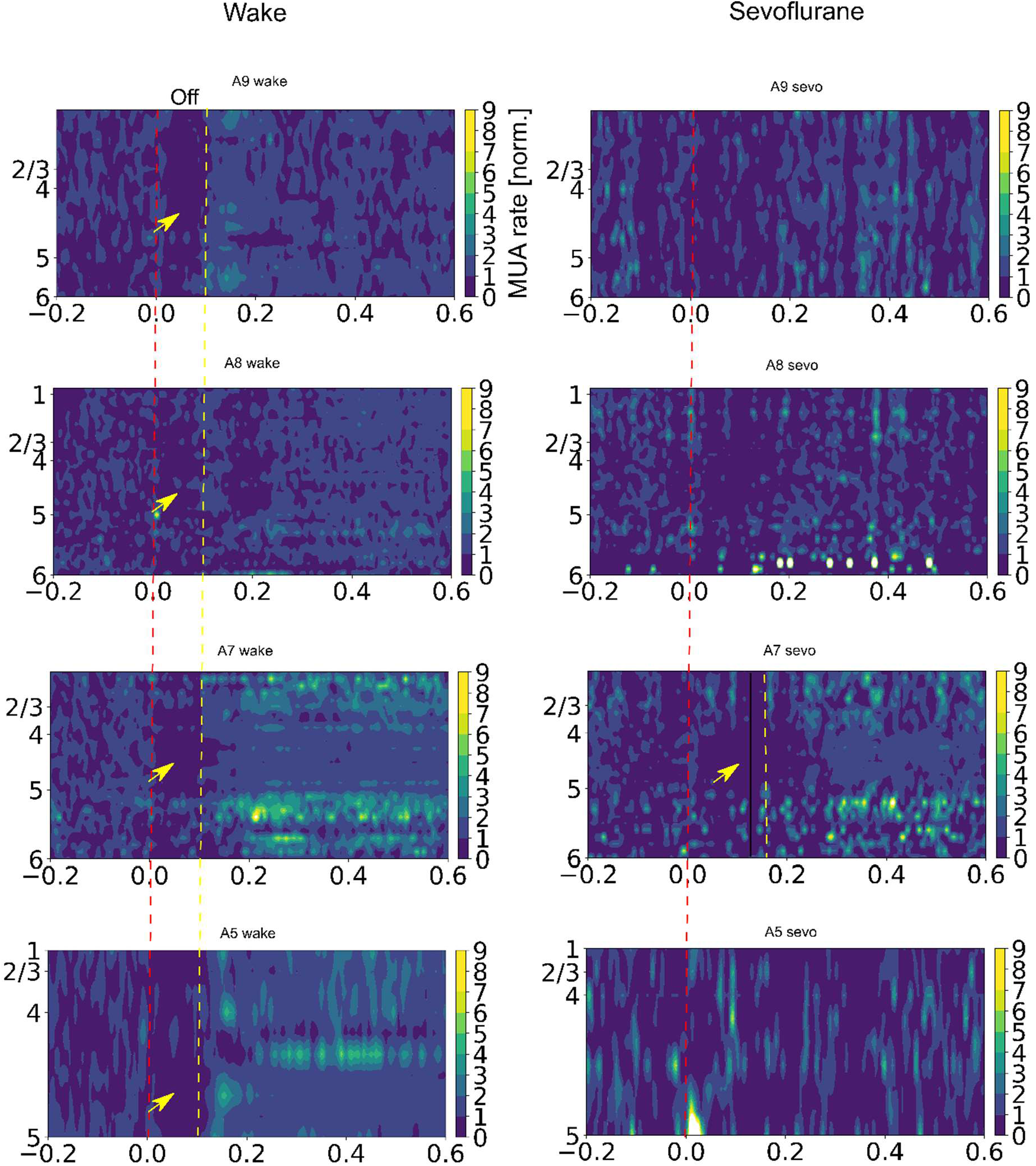
The off-period shows a higher variability during sevoflurane anesthesia compared to wakefulness across animals. Heat maps of mean MUA rates showing firing activity across all cortical layers following electrical stimulation (red dashed lines) in 4 mice (A9, A8, A7, A5) during wakefulness (left) and sevoflurane anesthesia (right). During anesthesia, a decrease in MUA rate (off-period) was observed in 1 mouse, in layer 5 (mouse A7, yellow arrow), but present in all mice during wakefulness. The off-period was identified by bootstrap statistic (see methods) which maintained only the significant decrease in MUA rates relative to baseline.

**Supplementary Fig. 2:**
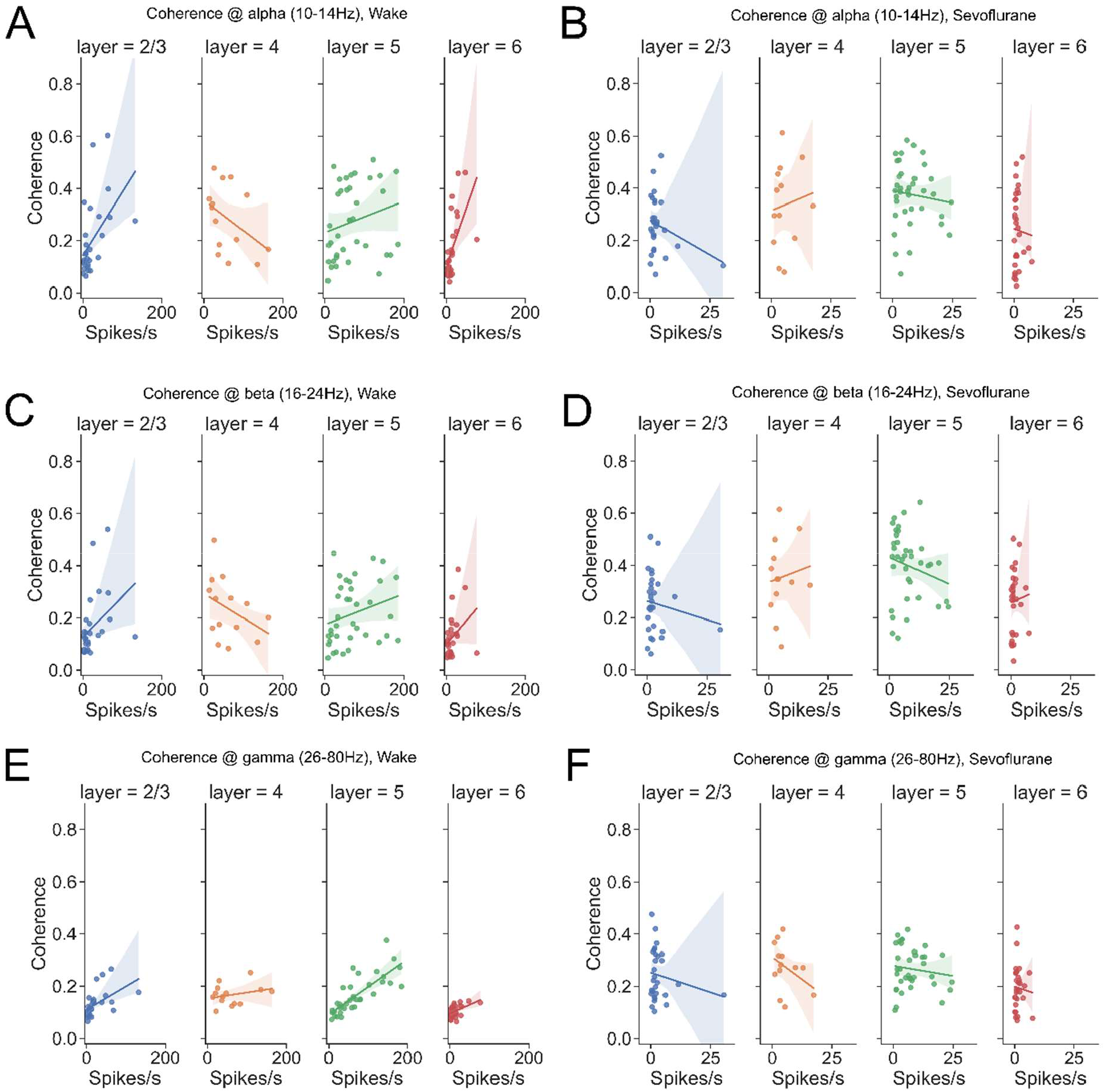
Spike-field coherence across cortical layers at alpha, beta and gamma frequencies. **A)** Spike-field coherence (wake) at alpha frequencies (∼10-14Hz) during late ERP (0.09 to 0.3s) as a linear function of spike rate across cortical layers (R-squared, Pearson coefficient, slope coefficient, P-value (pearson), P-value (slope_coeff)): l2/3 (0.25, 0.5, 0.0024, *0.01, *0.01), l4 (0.18, -0.43, -0.0012, 0.07, 0.07), l5 (0.05, 0.22, 6e-4, 0.1, 0.09), l6 (0.35, 0.59, 0.004,**0.002, **0.002). Permutation hypothesis test (Test statistic (pearson): pearson coefficient, H0: no correlation between rate and coherence; Test statistic (slope): slope coeffiecient of linear regression, H0: no linear relationship between rate and coherence, i.e. slope =0) **B)** Spike-field coherence (Sevoflurane) at alpha frequencies (∼10-14Hz) during late ERP (0.09 to 0.3s) as a linear function of spike rate across cortical layers (R-squared, Pearson coefficient, slope coefficient, P-value (pearson), P-value (slope_coeff)): l2/3 (0.08, -0.28, -0.005, 0.06, 0.06), l4 (0.014, 0.11, 0.004, 0.35, 0.35), l5 (0.01, -0.1, - 0.002, 0.27, 0.27), l6 (0.002, -0.04, -0.003, 0.43, 0.42). **C)** Spike-field coherence (wake) at beta frequencies (∼16-24Hz) during late ERP (0.09 to 0.3s) as a linear function of spike rate across cortical layers (R-squared, Pearson coefficient, slope coefficient, P-value (pearson), P-value (slope_coeff)): l2/3 (0.15, 0.39, 0.002, *0.047, *0.047), l4 (0.13, -0.36, 9e-4, 0.11,0.11), l5 (0.07, 0.26, 6e-4, 0.06, 0.06), l6 (0.12, 0.35, 0.002, 0.06, 0.06) **D)** Spike-field coherence (Sevoflurane) at beta frequencies (∼16-24Hz) during late ERP (0.09 to 0.3s) as a linear function of spike rate across cortical layers (R-squared, Pearson coefficient, slope coefficient, P-value (pearson), P-value (slope_coeff)): l2/3 (0.02, -0.14, -0.003, 0.24, 0.24), l4 (0.014, 0.12, 0.004, 0.35, 0.35), l5 (0.04, -0.21, - 0.004, 0.11, 0.11), l6 (0.004, 0.06, 0.004, 0.38, 0.38) **E)** Spike-field coherence (wake) at gamma frequencies (∼26-80Hz) during late ERP (0.09 to 0.3s) as a linear function of spike rate across cortical layers (R-squared, Pearson coefficient, slope coefficient, P-value (pearson), P-value (slope_coeff)): l2/3 (0.28, 0.53, 9e-4, **0.007, **0.007), l4 (0.15, 0.39, 3e-4, 0.11, 0.11), l5 (0.63, 0.79, 0.001, ***<1e-5, ***<1e-5), l6 (0.25, 0.5, 7e-4, **0.001, **0.001) **F)** Spike-field coherence (Sevoflurane) at gamma frequencies (∼26-80Hz) during late ERP (0.09 to 0.3s) as a linear function of spike rate across cortical layers (R-squared, Pearson coefficient, slope coefficient, P-value (pearson), P-value (slope_coeff)): l2/3 (0.03, -0.18, -0.003, 0.18, 0.18), l4 (0.14, -0.37, -0.007, 0.11, 0.11), l5 (0.02, -0.13, - 0.002, 0.2, 0.2), l6 (0.004, -0.06, -0.003, 0.39, 0.39)

## Supplementary material

### Methods

#### Experimental procedures

After recovery from surgery, we gradually habituated mice to the experimental environment, typically over the course of 3 days, starting with ∼5min on day 1 and increasing up to 25 min spent in the setup on day 3. In the beginning and end of each session, the mice received sweetened condensed milk as a reward. During this procedure, mice were placed in a transparent acrylic cylinder, leaving head and head bar exposed for fixation in the setup with a screw clamp. A heating pad was wrapped around the cylinder to maintain the mouse body temperature at ∼37-38°C during sevoflurane anesthesia. During wakefulness, the heating pad maintained a cylinder temperature of ∼25°C, which reduced muscle tone artefacts typically seen during EEG recordings at colder room temperatures.

#### Histology

We immersed the fixed brain in 15-30 % sucrose in PBS and stored it for ∼2 days at 4°C, typically at the time when the brain sank to the bottom of the flask. The brain was cut along the midline and both hemispheres were covered with OCT embedding medium (VWR) and quickly frozen with dry ice. 100µm thick sagittal sections were cut using a cryostat (NX70, Thermo Scientific), transferred to microscope slides and stained with Neurotrace™ green fluorescent nissl stain (Thermo Fisher, diluted 1:100-150 in 0.01M PBS). Slices were covered with Mowiwol™ (Sigma-Aldrich) mounting medium and stored at 4°C. Fluorescent images of brain sections were acquired with an AxioScan Z1 slide-scanning microscope (Carl Zeiss) at 10 x magnification and positions of electrodes were estimated using ZEN lite imaging software (Carl Zeiss). The average depth of implanted stimulation electrodes was 936µm ± 121 µm (std), n = 5 mice, (∼ layer 5). To determine positions of probe electrodes within the cortical layers we overlaid the fluorescent Nissl/DiI image with a reconstructed probe image and with sagittal Atlas/Nissl images of the Allen Mouse Brain Atlas (Lein, Hawrylycz et al. 2007).

#### Preprocessing of EEG, LFP and MUA

Raw EEG, LFP and MUA data were visually inspected, and channels accepted for further analysis in absence of large artefacts and low signal-to-noise ratio relative to other channels. LFP signals were pairwise referenced between channels of left and right probe shanks located at the same cortical depth. The stimulation artefact was removed using spline interpolation (1 ms before to 5 ms after stimulation) and EEG/ LFP data were band-pass filtered between 0.5 and 80 Hz using a 3^rd^ order butterworth filter applied in forward and backward direction using functions in Scipy 1.5.0. EEG/ LFP signals were down sampled to 500 Hz and the offset removed by subtracting the mean of the baseline period from each channel/trial. Trials with large artefacts were removed if larger than a threshold value defined by 3 x the standard deviation of the averaged root mean square (rms) of a baseline period (-1 to 0s) and a period following stimulation (0 to 1 s). MUA signals were referenced using the common median of all channels and the stimulation artefact was removed by linear interpolation ranging from 3 ms before to 4 ms after stimulation. MUA data were band-pass filtered between 300 and 6000 Hz using a 3^rd^ order butterworth filter applied in forward and backward direction.

#### Spontaneous cortical activity during wakefulness and anesthesia

The spontaneous cortical activity associated with the different experimental conditions was quantified from 86 epochs of 2 seconds of epidural EEG signal that preceded the electrical stimulation (from -2 to 0 s). A bipolar frontal-occipital derivation (M2-V2/RS right) was chosen for analyzing spontaneous activity (Arena, Lamanna et al. 2017, Arena, Comolatti et al. 2021) and a Morlet wavelet convolution (3 cycles, 40 wavelets, linearly spanning from 2 to 80 Hz) was performed on each epoch. Instantaneous spectral powers were normalized by 1 and converted in dB or were extracted and averaged across samples and trials, obtaining an estimation of the power of each frequency for each animal and condition. The resulting periodogram was linearly fitted in Log-Log coordinates, in the frequency range 20-40 Hz. The slope of the obtained linear function was the spectral exponent of the 1/f function and was used to quantify the distribution of frequency powers in the spontaneous EEG activity (Gao, Peterson et al. 2017, Colombo, Napolitani et al. 2019, Arena, Comolatti et al. 2021). Burst detection was performed for each epoch, on the instantaneous spectral powers averaged in range 5-40Hz. The average power activity across trials was assumed to follow a Rayleigh distribution. A statistical threshold was set at alpha = 0.05 and all the power increments from the averaged activity were extracted. Subsequent events that occurred within 300 ms were considered to be part of the same event and all the resulting events that lasted more than 50 ms were selected as potential bursts. The duration of all the bursts was summed and subtracted from the overall duration of spontaneous activity, estimating the time spent in suppression, with an EEG signal of very low amplitude. The anesthetized state was then quantified by the burst-suppression ratio (BSR), which specifies the fractional time spent in suppression (100%, all suppression; 0%, no suppression; (Vijn and Sneyd 1998, Arena, Lamanna et al. 2017)).

#### Power and phase coherence of EEG, LFP

We extracted spectral powers and phases of ERPs using Morlet wavelet convolution (3 cycles, 40 wavelets ranging from 2 to 80 Hz) as previously described (Arena, Comolatti et al. 2021). Powers of each channel/trial were normalized, i.e. divided by the averaged power across trials in a baseline window (-0.5 to -0.2 s), averaged across trials and converted to dB. To identify significant dB variations from baseline following stimulation, 95% confidence intervals (CI) of max/min dB values during baseline (-1 to -0.2 s) were calculated by resampling all trials with replacement (500 bootstrap replicates). The dB values above the 95% percentile were conserved, not significant values below set to 0. The phase coherence of ERPs across trials was calculated using inter-trial phase clustering (ITPC, (Cohen 2014, Arena, Comolatti et al. 2021, Arena, Juel et al. 2022)), for each, channel, wavelet frequency and time point and averaged across trials. ITPC ranges from 0 (no phase-locking) to 1 (maximum phase-locking). Only significant ITPC variations relative to baseline (-1 to -0.2 s) were conserved by calculating the 95% CI of max. ITPC values (500 bootstrap replicates). The ITPC values above the 95% percentile were conserved, not significant values below set to 0. The “ITPC drop time” was defined as the time point (in seconds) of the last significant ITPC value, averaged across a broadband frequency range of 6-80 Hz, in a time window of 800 ms following the stimulation. To quantify the phase coherence between different cortical areas and layers we calculated “inter site phase clustering” (ISPC) as previously described (Cohen 2014, Arena, Comolatti et al. 2021, Arena, Juel et al. 2022). ISPC represents the consistency of the phase difference between EEG/LFP signals of two channels across trials, at each time-frequency point. ISPC of each channel pair and frequency is then baseline corrected by subtracting the corresponding average value in the time window from -0.5 to -0.2 s. Bootstrap statistic (500 replicates) was performed to maintain only the significant ISPC variations from baseline, i.e. values above the 95% percentile of max/min ISPC values. All ISPC scores that could be explained by volume conduction (clustering of phase difference around 0 or pi) were excluded from the analysis by using the Gaussian v test (Cohen 2014, Arena, Comolatti et al. 2021). The ISPC scores < 0 were also excluded from analysis and set to 0, in order to consider only the positive ISPC scores, which represented the significant functional connections between channel pairs (same procedure of Arena et al. 2021). The resulting ISPC values from each channel pair were averaged in the frequency range 6–8 Hz and in the time window 0.09-0.3 s.

#### PCI^*st*^

PCI^*st*^ was used to estimate the capacity for consciousness (Comolatti, Pigorini et al. 2019, Arena, Comolatti et al. 2021), which quantifies the spatiotemporal complexity of mean EEG/LFP responses following cortical stimulation. It was assessed, from 0 to 600 ms following stimulation, and across time, in short, moving windows of 100 ms, with 20 ms of overlap. PCI^*st*^ was computed using open-access code (github.com/renzocom/PCIst) with the following parameters: ‘baseline_window’: (- 500,-5), ‘k’: 1.2, ‘min_snr’: 2.0, ‘max_var’: 99, ‘n_steps’:100. The 95% confidence intervals of difference in mean PCI^*st*^ values between wakefulness and anesthesia were calculated using 1000 bootstrap replicates.

#### MUA spike rate

Single spikes of MUA signals were detected when crossing a fixed threshold of -50 µV, corresponding likely to neurons with somata at an approximate distance of x < 100 µm to the extracellular recording site (Henze, Borhegyi et al. 2000, Pettersen and Einevoll 2008), but may vary depending on the e.g. biophysical properties of these cells and initiation site of the action potential. The timestamp of a negative spike peak was extracted and binarized (1 = spike, 0 = no spike) for all samples, trials and channels. To calculate normalized spike rates binarized spikes were binned/summed in 10 ms time windows with 5 ms overlap between adjacent windows. Spike rates were normalized to baseline firing rates, i.e. by dividing the spike rate/s for each bin and trial by the mean spike rates averaged across, bins and trials during baseline period. The “off-period” following stimulation (Fig. 4B) was defined as the mean duration (s) when spike rate dropped across all layers and animals. Bootstrap statistic (500 replicates) was performed to maintain only the significant spike rate variations from baseline, i.e. above the 95% percentile of minimum spike rates averaged across trials. The “on-period” represents the time window when spike rate increased again following the “off-period” (∼0.09 s). To investigate how MUA spike rates relate to phase-locked LFP responses, the end of the “on-period” was defined as the time point of maximum ITPC drop (at 6-8 Hz) across animals, averaged across trials and channels (∼0.3 s). The “on-period” window was used to calculate firing rates, spike-field coherence (Fig. 4), and functional connectivity (Fig.5) between cortical layers/areas during wakefulness and sevoflurane anesthesia.

#### Spike-field coherence

The coupling between the MUA and LFP signals was quantified during the “on-period” (0.09 to 0.3 s) following stimulation using spike-field coherence (Lepage, Kramer et al. 2011) implemented in Python (https://mark-kramer.github.io/Case-Studies-Python/11.html). In brief, the field spectra and cross-spectra of mean subtracted LFP and binarized spike data are computed for each channel and trial using the Fast Fourier Transform (FFT), and averaged across all trials:

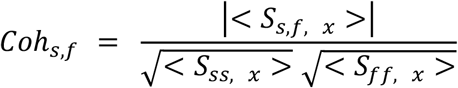

Where |< *S*_*s,f, x*_ >| indicates the magnitude of the trial averaged cross spectrum for each frequency x, and 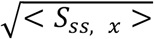 and 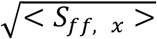 the square root of the magnitude of the trial averaged field spectra of *s* (spike) and *f* (field) time series. The resulting coherence values of each channel and frequency were then averaged across frequency bands: theta (6-8 Hz), alpha (10-14 Hz), beta (16-24 Hz) and gamma (26-80 Hz).

#### Statistics

Descriptive statistics in text and figure captions are given as mean ± one standard deviation, unless otherwise stated. Boxplots shown in figures were calculated using Seaborn and Matplotlib boxplot functions, and indicate median and upper/lower quartile values. Linear regression models were fitted using ordinary least squares method (Scikit-learn, Statsmodels, Seaborn) and slope, goodness of fit (R^2^) and Pearson’s correlation coefficient computed. To estimate 95% confidence intervals of mean and difference of means (Fig. 2G, H), bootstrap resampling was performed using numpy 1.19. P-values were obtained by bootstrap and permutation hypothesis tests as noted in the figure legends, hence no assumptions about the probability distribution underlying the data were assumed. To simulate p-values we defined first our hypothesis and calculated the observed test statistic of interest. We then computed bootstrap or permutation replicates (10^4^ -10^5^ samples) of our test statistic by simulating the null hypothesis. The probability was then calculated by determining the fraction of results at least as extreme as the observed test statistic (1-tailed), under the assumption that the null hypothesis is true. P-values in the figures and legends are marked with * if p < 0.05, ** if p < 0.01, *** if p < 0.001. P-values were corrected for multiple comparisons if necessary (see figure legends).

